# Lymphotoxin β receptor: A crucial role in innate and adaptive immune responses against *Toxoplasma* g*ondii*

**DOI:** 10.1101/2021.01.11.426315

**Authors:** Anne Tersteegen, Ursula R. Sorg, Richard Virgen-Slane, Marcel Helle, Patrick Petzsch, Ildiko R. Dunay, Karl Köhrer, Daniel Degrandi, Carl F. Ware, Klaus Pfeffer

## Abstract

The LTβR plays an essential role in the initiation of immune responses to intracellular pathogens. In mice, the LTβR is crucial for surviving acute toxoplasmosis, however, up to now a functional analysis is largely incomplete. Here, we demonstrate that the LTβR is a key regulator required for the intricate balance of adaptive immune responses. *T. gondii* infected LTβR^−/−^ mice show globally altered IFNγ regulation, reduced IFNγ-controlled host effector molecule expression, impaired T cell functionality and an absent anti-parasite specific IgG response resulting in a severe loss of immune control of the parasites. Reconstitution of LTβR^−/−^ mice with toxoplasma immune serum significantly prolongs the survival following *T. gondii* infection. Notably, analysis of RNAseq data clearly indicates a specific effect of *T. gondii* infection on the B cell response and isotype switching. This study unfolds the decisive role of the LTβR in cytokine regulation and adaptive immune responses to control *T. gondii*.

## Introduction

The lymphotoxin β receptor (LTβR) is one of the core members of the tumor necrosis factor (TNF)/TNF receptor (TNFR) superfamily (1, 2). It has two cognate ligands, LTβ (LTα_1_β_2_) and LIGHT (homologous to lymphotoxins, exhibits inducible expression, and competes with HSV glycoprotein D for herpes virus entry mediator [HVEM], a receptor expressed by T lymphocytes) (3, 4). LTβR mediated signaling is known to be essential for the organogenesis of secondary lymphoid tissues, the maintenance of their structure and its role in mediating innate immune responses to many pathogens is also well documented (2, 5–7). LTβR deficient (LTβR^−/−^) mice lack of lymph nodes (LNs) and Peyer’s patches (PPs) and show reduced numbers of natural killer (NK) and dendritic cells (DCs) as well as impaired immunoglobulin (Ig) affinity maturation (7, 8). In infection models, LTβR^−/−^ mice show pronounced defects in their immune response against *Listeria monocytogenes*, *Mycobacterium tuberculosis* (5), cytomegalovirus (9), LCMV (10) and Zika virus (11) as well as *Toxoplasma gondii* (*T. gondii*) (12). In spite of these extensive deficits, not much is known about the exact role of LTβR signaling for an efficient generation of the immune response against pathogens.

*T. gondii*, the causative agent of toxoplasmosis, is an obligate intracellular parasite belonging to the Apicomplexa. It is able to invade most warm-blooded vertebrates including humans (13, 14) and can infect all nucleated cells. While acute toxoplasmosis usually presents with only mild, flu-like symptoms in immunocompetent hosts, it sometimes manifests as lymphadenitis, hepatosplenomegaly, myocarditis or pneumonia. In immunocompromised patients toxoplasmosis can cause serious health problems and, when primary infection occurs during pregnancy, severe congenital defects may occur (15–17).

The early immune response to *T. gondii* is characterized by recognition of *T. gondii* associated molecules (i.a. profilin) by different cell types such as DCs. These cells produce distinct cytokines in response to infection such as IL-12 and TNF thus activating and stimulating other cell types including NK cells (18), T cells (19), ILCs (20), and macrophages (21) which in turn produce inflammatory cytokines such as IFNγ.

IFNγ signalling is essential for limiting *T. gondii* proliferation during the acute stage of toxoplasmosis and driving the parasite into the chronic stage where it is contained by a functional immune response (22–25). IFNγ driven effector mechanisms include induction of cell-autonomous effector mechanisms (26, 27), such as depletion of tryptophan (28) and reactive nitrogen production (29) which suppress *T. gondii* replication and are essential for restricting parasite growth. IFNγ also strongly induces murine Guanylate-Binding Proteins (mGBPs) which play a major role in restricting parasite growth of *T. gondii* as well as other intracellular pathogens (30–33). Within an infected cell, *T. gondii* resides within a parasitophorous vacuole (PV) that effectively protects the parasite from lysosomal activity (34). mGBPs are recruited to the PV and are instrumental in destroying first the PV and then the parasites within (30, 31, 33, 35, 36).

Previous studies have shown that other core members of the TNF/TNFR superfamily such as the ligands TNF, LTα which signal via the TNFRI receptor also play an important part in the immune response to *T. gondii* (25, 37, 38). However, there is only limited data published on the role of the LTβR: It has been demonstrated that signaling via the LTβR is essential for the up-regulation of mGBPs after *T. gondii* infection as well as for overall survival (12). Glatman Zaretsky et al. have shown that LTβ signaling is important for maintaining intact splenic architecture and, indirectly, for efficient *T. gondii* specific antibody production (39). Nevertheless, the pathophysiology responsible for the increased susceptibility of LTβR^−/−^ mice to *T. gondii* infection is still elusive.

Here, we demonstrate that LTβR deficiency results in dramatically dysregulated IFNγ responses, impaired expression of anti-parasite effector molecules, limited T cell functionality and an abrogated *T. gondii* specific IgG response. We show that by transfer of *T. gondii* immune serum survival of LTβR^−/−^ mice can be prolonged, demonstrating that the susceptibility of LTβR^−/−^ mice to *T. gondii* infection is possibly due to a direct role of LTβR signaling in Ig class switch. These results lead to a new understanding of LTβR mediated immunity and the pathophysiology of toxoplasmosis and will hopefully aid in developing much needed new treatment and prevention options such as passive vaccination strategies for human toxoplasmosis.

## Results

### LTβR deficiency leads to increased parasite burden in lung, spleen and muscle

While wildtype C57BL/6 (WT) mice survive a *T. gondii* infection, LTβR^−/−^ mice are highly susceptible to *T. gondii* infection and do not survive beyond day 14 *p.i.* (Fig. 1a). This high susceptibility is in accordance with our previous study (12). To characterize the cause of this susceptibility in LTβR^−/−^ mice we first assessed the parasite burden in *T. gondii* infected WT and LTβR^−/−^ animals during the acute phase of infection via qRT-PCR (Fig. 1b). In lung tissue, we found increasing amounts of *T. gondii* DNA up to day 10 *p.i.* in both cohorts with significantly higher amounts in LTβR^−/−^ compared to WT mice on day 10 *p.i.* In the spleen *T. gondii* DNA amounts increased only moderately in WT mice through the course of infection (Fig. 1b). In contrast, LTβR^−/−^ mice showed a significant increase of *T. gondii* DNA by day 10 *p.i.* and also significantly increased amounts compared to WT mice on days 7 and 10 *p.i.* Interestingly, in both genotypes reduced amounts of *T. gondii* DNA could be detected on day 10 compared to day 7 *p.i.* Similar results were observed in muscle tissue (Fig. 1b). In WT mice, the parasite burden rose only moderately, while LTβR^−/−^ mice showed a significant increase by day 10 *p.i.* as well as significantly higher amounts on days 7 and 10 *p.i.* compared to WT mice. To summarize, LTβR^−/−^ mice showed increased parasite burden compared to WT mice pointing towards a failure of these animals to adequately control parasite proliferation in the acute phase of infection.

**Fig. 1.**
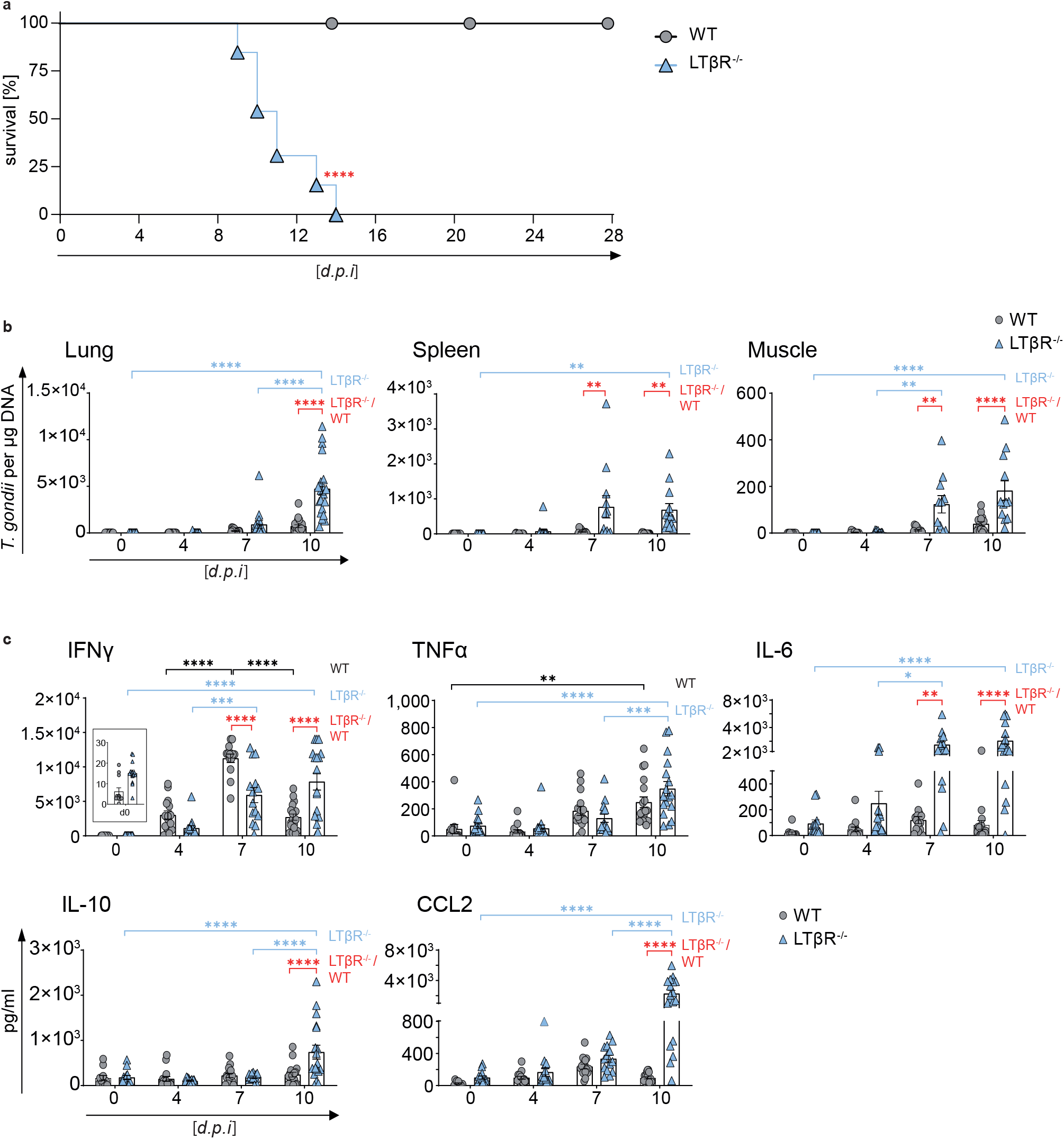
LTβR^−/−^ mice show increased parasite load and dysregulated cytokine expression. **a,** survival of *T. gondii* infected (ME49, 40 cysts, *i.p.*) WT (n=15) and LTβR^−/−^ (n=13) mice.**b**, qRT-PCR analysis of *T. gondii* DNA (assessing parasite load) in lung, spleen and muscle tissue of uninfected (d0) and *T. gondii* infected WT and LTβR^−/−^ mice (d0 - 7: n≥12, d10: n≥14). Expression of **c**, IFNγ, TNFα, IL-6, IL-10 and CCL2 in the serum of uninfected and *T. gondii* infected WT and LTβR^−/−^ mice (d0 - 7: n≥12, d10: n=18) analyzed via bead-based immunoassay. Data shown represent at least three independent experiments; symbols represent individual animals, columns represent mean values and error bars represent ± SEM. A log rank (Mantel Cox) test was used for statistical analysis represented in **a**. 2way ANOVA corrected for multiple comparison by the Tukey‘s post hoc test was used for statistical analysis represented in **b** and **c**. *P<0.0332, **P<0.0021, ***P<0.0002 and ****P<0.0001.

### Dysregulated cytokines in the serum of LTβR-/- mice after infection with *T. gondii*

Since cytokines, especially IFNγ and TNFα as signature molecules of a Th1 response play an important role in containing *T. gondii* expansion (16, 22, 40), we analyzed cytokine amounts in sera of infected mice (Fig. 1c). In both genotypes IFNγ amounts increased slightly by day 4 *p.i.* In WT animals, IFNγ amounts increased significantly by day 7 *p.i.* but was found to be markedly decreased again on day 10 *p.i.* While LTβR^−/−^ mice also showed a significant increase of IFNγ expression on day 7 *p.i.*, amounts were significantly lower than those of WT animals. Also, in LTβR^−/−^ mice IFNγ expression levels were significantly higher on day 10 *p.i.* compared to WT animals. TNFα expression increased significantly in WT as well as LTβR^−/−^ animals by day 10 *p.i.* and did not differ significantly between the two genotypes, although amounts in LTβR^−/−^ mice seemed to rise more steeply later in infection (day 7 vs. day 10 *p.i.* for WT and LTβR^−/−^ mice, respectively).

In WT animals expression of IL-6, another proinflammatory cytokine (41), was slightly increased on day 4 and day 7 *p.i.* but was reduced again on day 10 *p.i.* (Fig. 1c). In contrast, in LTβR^−/−^ mice IL-6 amounts rose significantly during the course of infection and were significantly higher on days 7 and 10 *p.i.* compared to WT mice. Amounts of IL-10, known for its anti-inflammatory properties during infection (42), did not change significantly in WT animals during the course of infection (Fig. 1c). In contrast, amounts in LTβR^−/−^ animals rose significantly on day 10 *p.i.* and were significantly higher compared to WT mice. The monocyte chemotactic factor (CCL2), a chemokine described to be induced by *T. gondii* (43), increased in WT as well as LTβR^−/−^ mice on days 4 and 7 *p.i.* But while CCL2 in WT mice declined again by day 10 *p.i.*, CCL2 further increased in LTβR^−/−^ mice on day 10 *p.i.* and were significantly higher than in WT mice (Fig. 1c). Interestingly, LTβR^−/−^ mice showed increased baseline amounts (day 0) for IFNγ, TNFα, IL-6, and CCL2 compared to WT mice, even though these differences were not significant.

Significantly different amounts were detected for IFNβ, IL-1α, IL-23, and IL-27 only on day 4 *p.i.* (Suppl. Fig. 1). LTβR^−/−^ animals showed increased baseline amounts (d0) for IFNβ, IL-1α, IL-1β, IL-17A, IL-23, IL-27, and IL-12p70, which were however significant only in the case of IL-1β. No differences in IL-12p70 levels could be detected for the two genotypes (Suppl. Fig. 1).

To summarize, uninfected LTβR^−/−^ mice show different baseline amounts of proinflammatory cytokines suggesting a subtle activation of the immune system. Furthermore, in these animals the coordinated immune defense during *T. gondii* infection is dysregulated.

### Markedly altered transcriptome in the lungs of LTβR^−/−^ mice after *T. gondii* infection

The lungs are one of the target organs of *T. gondii* tachyzoite dissemination (12, 44). In line with that observation, we detected high amounts of *T. gondii* DNA in lung tissue of LTβR^−/−^ compared to WT mice on day 4 *p.i.* (Fig. 1b). To determine whether WT and LTβR^−/−^ mice show differences in global gene expression patterns in the lungs, we analyzed lung tissue via RNAseq on day 7 *p.i.* Interestingly, gene set enrichment analysis (GSEA) of these data showed a significant upregulation of GO (biological process) molecular signatures for ‘response to type i interferons, response to interferon gamma, and interferon gamma mediated signaling pathway’ in *T. gondii* infected WT compared to LTβR^−/−^ mice on day 7 *p.i.* (Suppl. Fig. 2). The data depicted by a volcano plot (Fig. 2a) clearly shows a significant upregulation of IFNγ-regulated genes in *T. gondii* infected WT mice as compared to LTβR^−/−^ mice (day 7 *p.i*): For instance, transcripts for mGBPs (mGBP2b/1, 2, 6, 7, and 10), transcripts for effector molecules (IDO1, Gzmk), transcripts for chemokines and chemokine receptors responsible for recruitment of immune cells (CCL2, CCL4, CCL7, CXCL9, CXCL10, CCR1), transcripts for proteins involved in IFNγ-signaling (IRF1, STAT1), transcripts induced by IFNγ (TGTP1, PIM1) and other transcripts known to be involved in immune responses (CD274, IL12Rb1, Ly6i, Ly6c2, MMP8, RNF19b) were found to be highly expressed in infected WT, but not in LTβR^−/−^ lungs on day 7 *p.i.* This suggests that LTβR^−/−^ mice fail to adequately upregulate (IFNγ-dependent) immune responses in the lungs.

**Fig. 2.**
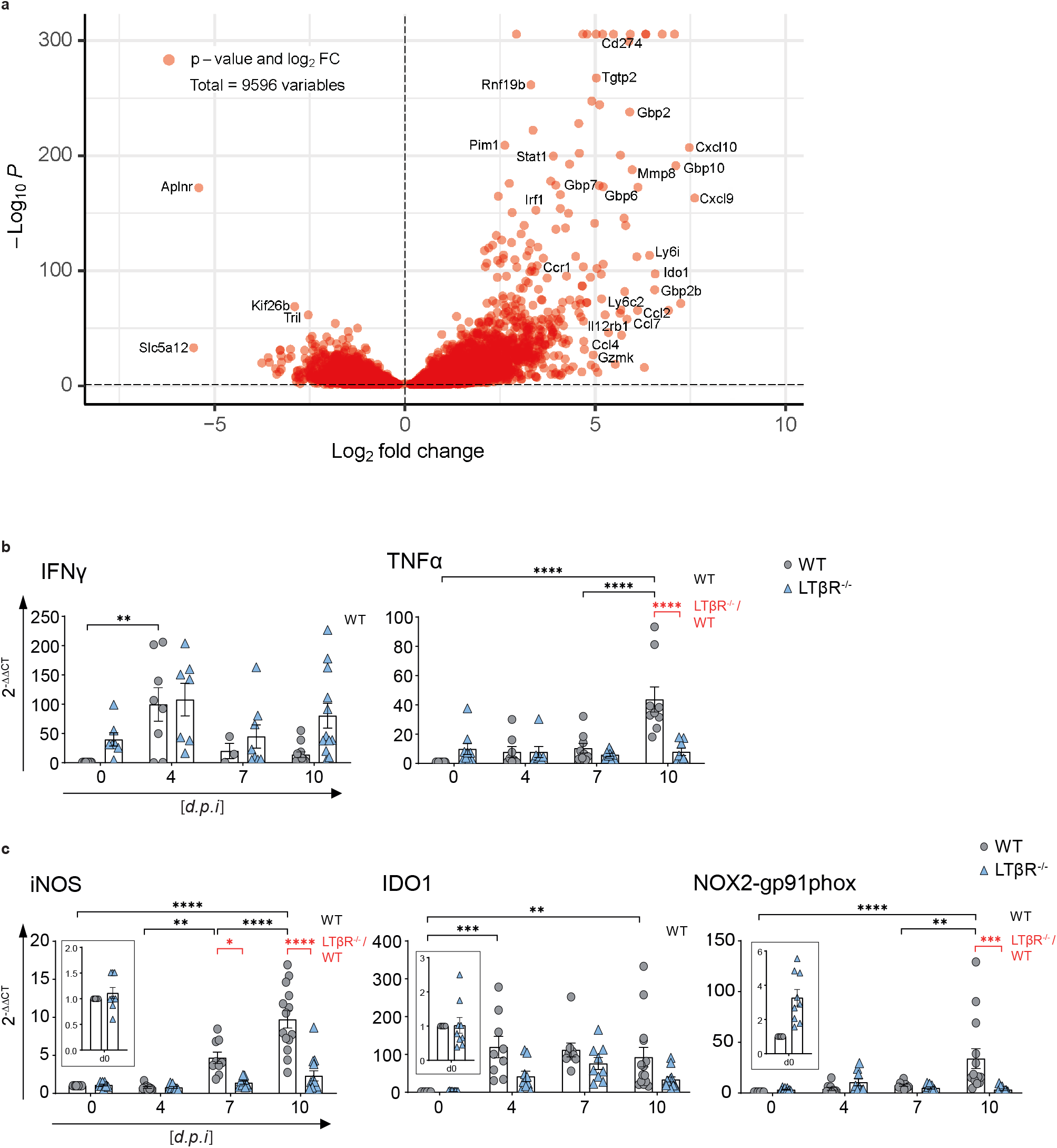
Lungs of LTβR^−/−^ mice show altered transcriptome after *T. gondii* infection. **a**, volcano plot showing RNAseq data of lung tissue of infected WT mice correlated to infected LTβR^−/−^ mice (d7 *p.i.*; n=3/group). Dashed horizontal black line represents an adjusted p-value of 0.1 (“Wald” test). qRT-PCR analysis of **b**, cytokines (IFNγ and TNFα) and **c**, host effector molecules (iNOS, IDO1, NOX2-gp91phox) in lung tissue from uninfected (d0) and *T. gondii* infected (ME49, 40 cysts *i.p.*) WT and LTβR^−/−^ mice (d0 - 7: n≥12, d10: n≥14; exception: IFNγ n≥3, d0 – 10 *p.i.*). Data shown in **b** & **c** represent four independent experiments; symbols represent individual animals, columns represent mean values and error bars represent ± SEM. 2way ANOVA corrected for multiple comparison by the Tukey‘s post hoc test was used for statistical analysis. *P<0.0332, **P<0.0021, ***P<0.0002 and ****P<0.0001.

### LTβR deficiency leads to dysregulation of cytokine expression in the lung

To extent RNAseq data (Fig. 2a) and cytokine levels in serum (Fig. 1c & Suppl. Fig. 1), we determined mRNA expression levels of cytokines in the lungs of infected WT and LTβR^−/−^ mice at several time points after infection (Fig. 2b & 2c). Baseline expression levels of IFNγ were higher in LTβR^−/−^ animals, thus, while levels rose on day 4 *p.i.* in both genotypes, this increase was only significant in WT mice (Fig. 2b). While IFNγ mRNA levels were markedly decreased in WT mice by day 10 *p.i.*, they were still markedly but not significantly elevated in LTβR^−/−^ mice (Fig. 2b). Baseline expression of TNFα was increased in LTβR^−/−^ mice but did not change significantly during the course of infection. WT mice showed a significance increase in TNFα expression on day 10 *p.i.* (Fig. 2b) indicating a significant difference in the cytokine response between the two genotypes on day 10 *p.i.* Baseline expression levels of LTβ were significantly increased in LTβR^−/−^ mice, which could be due to a lack of negative feedback or compensatory mechanisms. However, while levels tended to be higher in LTβR^−/−^ animals throughout the infection, there were no significant differences in LTβ expression between the two genotypes (Suppl. Fig. 3). IL-4 expression was significantly increased in WT animals on day 10 *p.i.* compared to baseline expression. In LTβR^−/−^ mice IL-4 expression was comparable to those of WT mice but not significantly increased on day 10 *p.i.* compared to baseline expression (Suppl. Fig. 3). These data confirm that LTβR^−/−^ mice show a dysregulated immune homeostasis not only in serum (Fig. 1c & Suppl. Fig.1) but also in lung tissue after *T. gondii* infection.

### LTβR deficiency leads to impaired IFNγ-regulated effector molecule expression in the lung

IFNγ-regulated effector molecules are pivotal in *T. gondii* elimination (31, 33, 45), having important immune response functions. In particular, the roles of effector molecules, such as iNOS, IDO, and NOX2-gp91phox (46–48) are well-documented. Since RNAseq data (Fig. 2a) showed high expression of effector molecules in infected WT, but not LTβR^−/−^ (31) mice we assessed the expression of major effector molecules in lungs by qRT-PCR next (Fig. 2c). In contrast to WT mice, LTβR^−/−^ mice failed to up-regulate iNOS expression post infection leading to significant differences between the two genotypes on day 7 and 10 *p.i.* WT mice showed significant upregulation of IDO1 expression on day 4 *p.i.* and had significantly increased IDO1 expression levels on day 10 *p.i*, whereas LTβR^−/−^ mice showed only a minor increase of IDO1 expression and this difference was not significant compared to baseline expression. NOX2-gp91phox presented a similar picture: Significantly increased NOX2-gp91phox expression in WT animals on day 10 *p.i.* compared to baseline expression as well as compared to LTβR^−/−^ mice and a complete failure of upregulation of NOX2-gp91phox in the absence of LTβR. The failure to adequately upregulate IFNγ-regulated effector molecules involved in cell intrinsic defense mechanisms essential for suppressing *T. gondii* replication most likely contributes to the increased parasite burden observed in LTβR^−/−^ animals.

### LTβR deficiency leads to impaired IFNγ-induced mGBP expression and IFNγ signaling in the lung

Another important group of genes upregulated in an IFNγ-dependent manner after *T. gondii* infection are mGBPs (30). These GTPases have been shown to be essentially involved in *T. gondii* elimination (30–33). A heat map for mGBP expression data (Fig. 3a) was generated from the RNAseq data illustrating an overall slight increase in baseline mGBP expression in uninfected (day 0) LTβR^−/−^ mice compared to WT mice but an overall lower mGBP expression in LTβR^−/−^ compared to WT mice on day 7 *p.i.* These results were confirmed by qRT-PCR analysis of mGBP mRNA expression: For all mGBPs analyzed (mGBP1, 2, 3, 5, 6/10, 7, 8 and 9) we observed a significant increase in mGBP expression (P<0.0001 in all cases) in WT animals by day 10 *p.i.* (Fig. 3b). In contrast, in LTβR^−/−^ mice a significant rise on day 10 *p.i.* compared to baseline expression was only observed for mGBP2, mGBP3 and mGBP7. Also, expression levels of all mGBPs were significantly higher in WT mice compared to LTβR^−/−^ mice on day 10 *p.i.* with the exception of mGBP6/10 where expression levels were only subtly increased in WT mice. The failure to adequately upregulate expression of mGBPs early after *T. gondii* infection was further confirmed by immunoblot analysis, where upregulation of mGBP2 and mGBP7 protein expression was already detectable on day 4 *p.i.* in WT mice but not in LTβR^−/−^ mice (Fig. 3c and Suppl. Fig. 4). This defect in upregulation of mGBP expression after *T. gondii* expression likely has a major effect on the ability of LTβR^−/−^ mice to contain parasite replication, as mGBPs are essential for an effective immune response against this parasite (31–33).

**Fig. 3.**
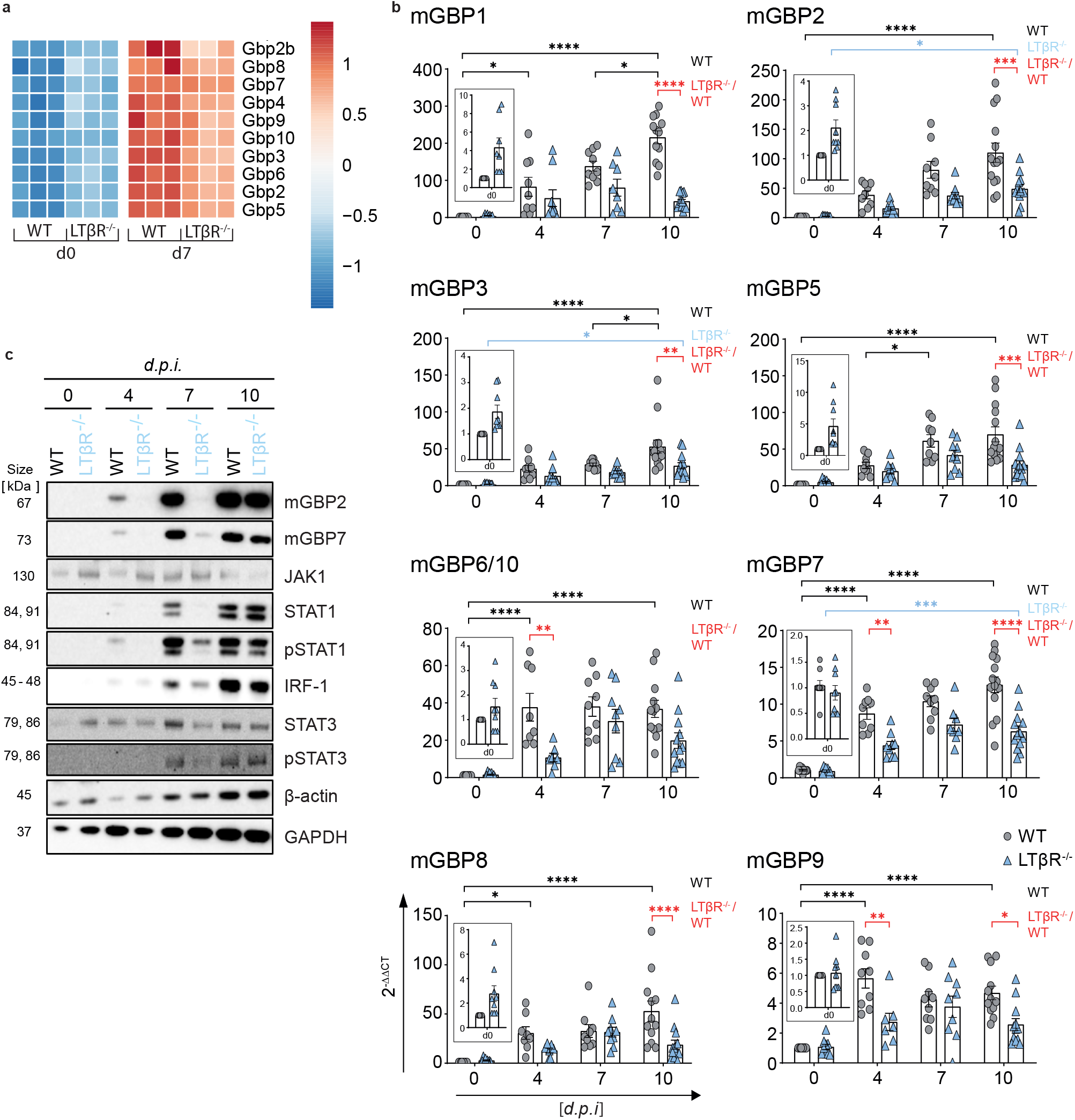
LTβR deficiency dysregulated IFNγ signaling in the lung. **a**, Heat map of differentially expressed murine guanylate-binding proteins (mGBPs) based on RNAseq analysis (“Wald” test & adjusted p-value of 0.1) of lung tissue from uninfected (d0) and *T. gondii* infected (ME49, 40 cysts *i.p.*, d7 *p.i.*) WT and LTβR^−/−^ mice (n=3). **b**, qRT-PCR of mGBPs in lung tissue from uninfected and *T. gondii* infected WT and LTβR^−/−^ mice (d0 - 7: n≥12, d10: n≥14). Data shown represent four independent experiments; symbols represent individual animals, columns represent mean values and error bars represent ± SEM. **c**, Immunoblot analysis of proteins involved in or induced via the IFNγ signaling pathway in lung tissue from uninfected and *T. gondii* infected WT and LTβR^−/−^ mice. 2way ANOVA corrected for multiple comparison by the Tukey‘s post hoc test was used for statistical analysis represented in **b**. *P<0.0332, **P<0.0021, ***P<0.0002 and ****P<0.0001. Data shown in **c** is a representative of three independent experiments.

Since protein expression of IFNγ-induced mGBPs was affected in lungs of LTβR^−/−^ mice in *T. gondii* infection we further analyzed protein expression of prototype genes directly involved in IFNγR signaling (Fig. 3c & Suppl. Fig 4). Protein expression levels of STAT1, pSTAT1, IRF-1, and pSTAT3 increased in WT mice during the course of infection. In contrast, LTβR^−/−^ animals showed a marked delay in the upregulation of these proteins. In WT animals, JAK1 and STAT3 expression increased until day 7 *p.i.* but decreased again on day 10 *p.i.* In uninfected LTβR^−/−^ mice, expression of these proteins was higher than in uninfected WT mice but did not increase early in infection. This also provides evidence for an altered IFNγ/IFNγR signaling axis during *T. gondii* infection.

To summarize, mRNA and protein expression data from the lungs indicate that uninfected LTβR^−/−^ animals show an activated immune status compared to WT animals, but fail to adequately upregulate IFNγ-dependent immune effector responses after *T. gondii* infection, possibly explaining the increased parasite burden and the subsequently increased infection susceptibility of LTβR^−/−^ mice.

### mGBP upregulation and recruitment to the PV after IFNγ stimulation *in vitro*

Since upregulation of mGBP expression was impaired in LTβR^−/−^ mice after *T. gondii* infection (Fig. 3), we asked whether IFNγ-dependent upregulation of mGBP expression was directly dependent on LTβR signaling (Fig. 4). We therefore analyzed whether LTβR^−/−^ mouse embryonic fibroblasts (MEFs) were able to upregulate mGBPs after IFNγ stimulation and whether mGBPs could recruit to the PV in infected, IFNγ pre-treated LTβR^−/−^ MEFs. After pre-incubation with IFNγ *in vitro*, *T. gondii* infected LTβR^−/−^ and WT MEFs showed comparable upregulation of all tested mGBPs (mGBP1, 2, 3, 5, 6/10, 7, and 9) with the exception of mGBP8 where WT mice showed increased mRNA expression (Fig. 4a). Also, after pre-incubation with IFNγ mGBP2 was able to recruit to the PV of *T. gondii* in LTβR^−/−^ MEFs (Fig. 4b). These results demonstrate that expression of mGBPs can be successfully induced in LTβR^−/−^ MEFs in the presence of exogenous IFNγ and that the lack of LTβR signaling appears not to interfere with the ability of mGBP2 to recruit to the PV in LTβR^−/−^ MEFs. This suggests, that the absence of LTβR signals do not impact IFNγR signaling required for mGBP function.

**Fig. 4.**
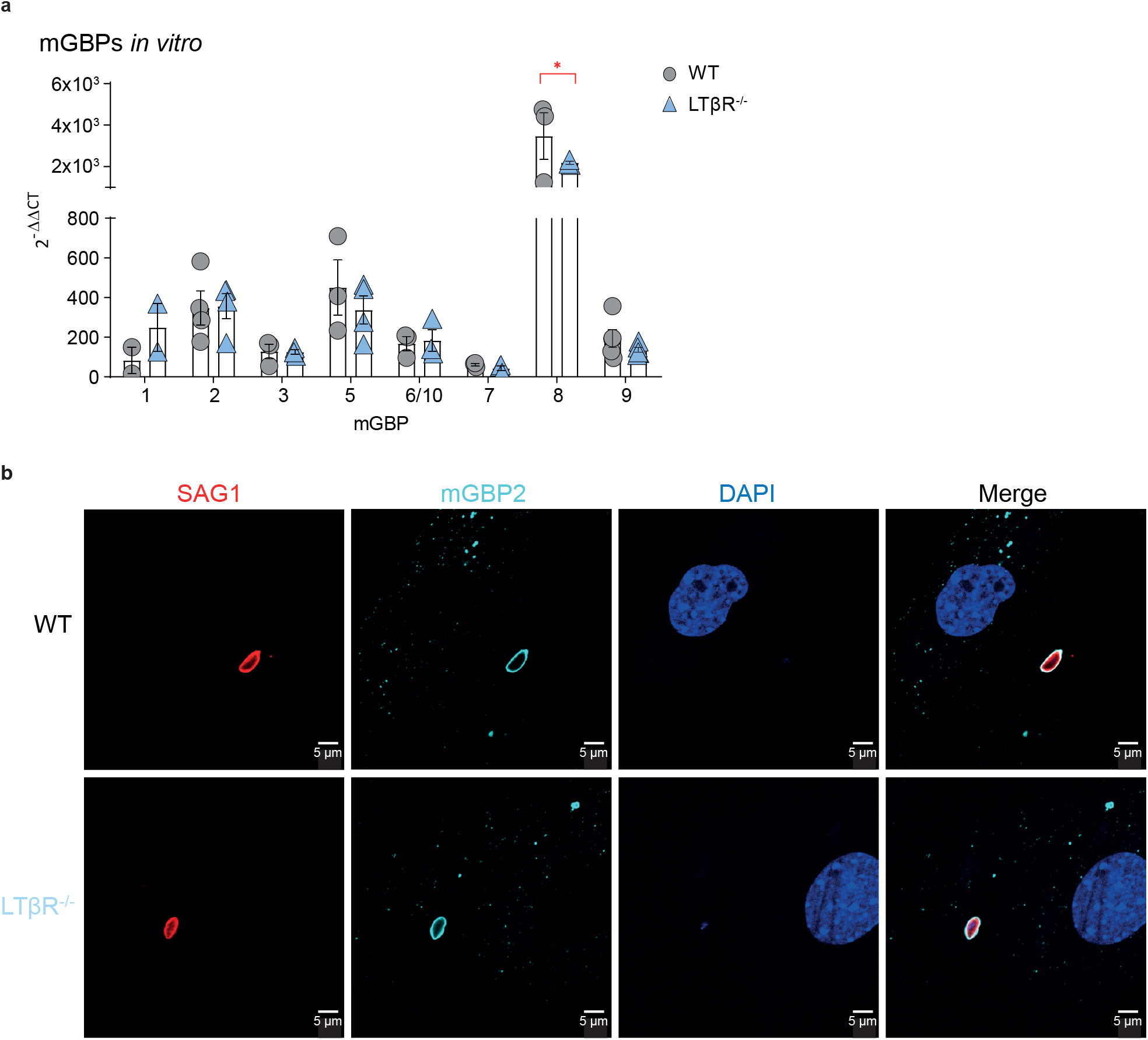
mGBP upregulation and recruitment. **a**, qRT-PCR analysis of mGBP mRNAs expression of uninfected WT and LTβR^−/−^ MEFs stimulated with IFNγ (7.5 ng/ml) for 8h (all n=3, except for mGBP1 where n=2). Each symbol represents an individual techniqual replicate; columns represent mean values and error bars represent ± SEM. 2way ANOVA corrected for multiple comparisons by the Sidak post hoc test was used for statistical analysis. *P<0.00332. **b**, Representative immunofluorescence analysis of *T. gondii* tachyzoite (MOI 1:40) infected WT and LTβR^−/−^ MEFs. Cells were prestimulated with IFNγ [7.5 ng/ml] for 16h before infected with *T. gondii* tachyzoites for 2h. *T. gondii* surface antigen SAG1 was visualized using a Cy3-conjugated and mGBP2 using an Alexa Fluor 633-conjugated secondary antibody for detection of mGBP2 recruitment towards the *T. gondii* PV. Cell nuclei were stained using DAPI. Data shown in **a** & **b** represent at least two independent experiments.

### Differences in spleen size and weight in LTβR^−/−^ mice

Since LTβR^−/−^ mice lack lymph nodes (7), the spleen is the primary organ where the immune response against *T. gondii* is primed. It has been described that during the acute phase of *T. gondii* infection the splenic architecture is disrupted transiently (39). When we compared spleens of WT vs. LTβR^−/−^ mice, spleens of the latter were markedly larger (Suppl. Fig. 5a) in uninfected (day 0) healthy animals. While spleens of both genotypes significantly increased in weight during the course of *T. gondii* infection, spleen weights of WT mice were significantly higher compared to LTβR^−/−^ mice on day 10 *p.i.* (Suppl. Fig. 5b). This increase of spleen weight in WT mice was not due to increased cellularity, as splenocyte counts were consistently higher in LTβR^−/−^ spleens before as well as on days 4 and 7 post infection (Suppl. Fig. 5c). By day 10 *p.i.*, cell numbers in the spleens of both genotypes were comparable, mostly due to a significant drop of splenocyte numbers in LTβR^−/−^ mice. This also indicates that the initial immune response in spleens of LTβR^−/−^ mice is disturbed.

### No apparent difference in T cell subpopulations in spleens of LTβR^−/−^ mice

Consecutively, we analyzed the composition of the splenocytes using flow cytometry (Fig. 5). Since T cells are essential to control *T. gondii* infection (49, 50), we analyzed T cell subpopulations in LTβR^−/−^ spleens (Fig. 5a). Analysis of absolute numbers of CD3^+^, CD4^+^, CD8^+^, activated (CD3^+^CD25^+^) T cells and *T. gondii* specific (pentamer^+^) CD8^+^ T cells (Fig. 5a) revealed almost no significant differences between WT and LTβR^−/−^ mice either before or during infection. The only exception were CD4^+^ T cells on day 4 *p.i.* where WT and LTβR^−/−^ mice showed a moderate decrease and increase, respectively. In both genotypes, numbers of activated CD3^+^CD25^+^ T cells were significantly increased on day 7 *p.i.* but LTβR^−/−^ mice showed similar numbers of total T cells. In addition, LTβR^−/−^ mice showed a comparable rise of activated CD3^+^CD25^+^ T cells on day 7 *p.i.* and a comparable expansion of *T. gondii* specific (pentamer^+^) CD8^+^ T cells on day 10 *p.i.* (Fig. 5a).

**Fig. 5.**
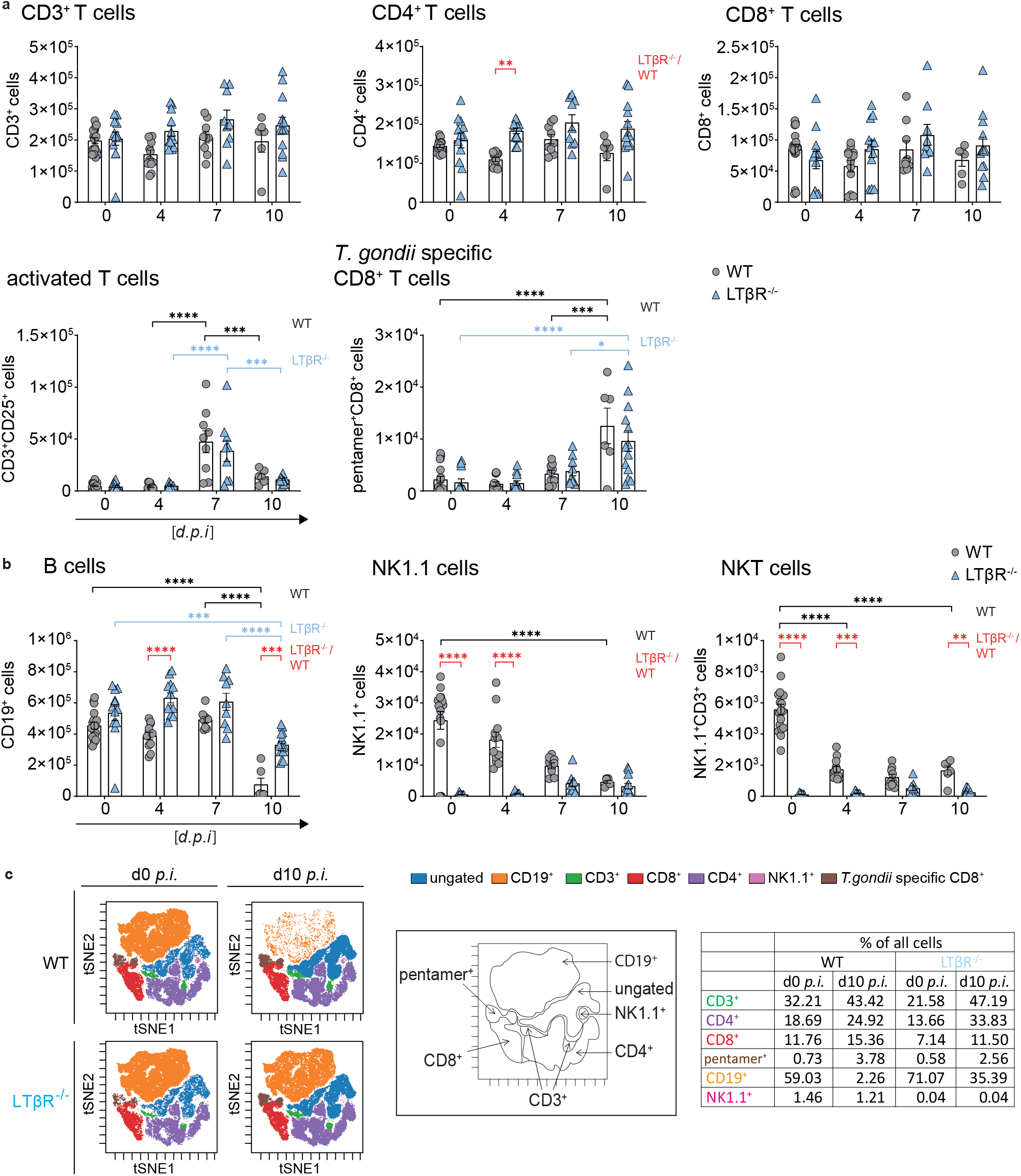
Dysregulated immune cell numbers in LTβR^−/−^ mice. **a**, Absolute cell numbers of CD3^+^, CD4^+^, CD8^+^, CD25^+^CD3^+^ and pentamer^+^CD8^+^ T cells and **b**, CD19^+^, NK1.1^+^ and NK1.1^+^CD3^+^ cells in spleens of uninfected (d0) and *T. gondii* infected (ME49, 40 cysts, *i.p.*) WT and LTβR^−/−^ mice (d0 – 7 *p.i.*: n=12, d10 *p.i.*: n≥6 determined via flow cytometry. **c**, Representative tSNE plots from splenocytes of uninfected and *T. gondii* infected (d10 *p.i.*) WT and LTβR^−/−^ mice. Clustered populations were identified using the indicated markers. Data shown represent at least three independent experiments; symbols represent individual animals, columns represent mean values and error bars represent ± SEM. 2way ANOVA corrected for multiple comparison by the Tukey‘s post hoc test was used for statistical analysis represented in **a** and **b**. *P<0.0332, **P<0.0021, ***P<0.0002 and ****P<0.0001.

Baseline numbers of CD19^+^ B cells were somewhat higher in LTβR^−/−^ mice and significantly increased on day 4 *p.i.*, but while numbers of CD19^+^ B cells dropped significantly in both genotypes on day 10 *p.i.,* they were still significantly higher in LTβR^−/−^ mice (Fig. 5b).

Since LTβR^−/−^ mice are known to have fewer NK and NKT cells (8, 51, 52), it was not surprising to observe that absolute NK1.1^+^ cells were significantly higher in WT compared to LTβR^−/−^ mice before infection and on days 4 and 7 *p.i.* (Fig. 5b). On day 10 *p.i.* NK1.1^+^ cell numbers of both genotypes were similar, due to the drop of NK1.1^+^ cells in spleens of WT mice during the course of infection. Similarly, NK1.1^+^CD3^+^ NKT cells in WT mice whose absolute numbers declined during the course of infection but were higher than those of LTβR^−/−^ mice before infection and on days 4 and 7 *p.i.* which is in accordance with published data (51). Unbiased analysis of the cytometry data set using tSNE (Fig. 5c) confirmed these data, notably the absence of NK1.1^+^ cells in uninfected LTβR^−/−^ mice (1.46 % and 0.04 %, respectively) and the marked drop in absolute CD19^+^ B cell numbers in WT mice by day 10 *p.i.* which was absent in LTβR^−/−^ animals (59.03 % to 2,26 % vs. 71.07 % to 35.39 %, respectively; Fig. 5c). This demonstrates that the deficiency of the LTβR does not impact T cell numbers after *T. gondii* infection, especially the expansion of parasite specific T cells, while it does seem to influence B cell numbers during the acute phase of *T. gondii* infection.

In conclusion, LTβR^−/−^ compared to WT mice do not show a significant difference in overall and antigen specific T cell numbers either before or after *T. gondii* infection, but B cell, NK1.1^+^ and NKT cell numbers appear to be significantly affected by the absence of LTβR before and during infection.

### Impaired T cell effector function in the spleen in the absence of the LTβR

Even though LTβR^−/−^ mice are highly susceptible to *T. gondii* infection we detected comparable CD8^+^ and *T. gondii* specific CD8^+^ T cell numbers in the spleen (Fig. 5a). We therefore decided to determine whether these T cells were fully differentiated and functional with regard to their ability to produce IFNγ, contained cytotoxic granules (GzmB^+^ and perforin^+^) and were able to degranulate (CD107a^+^ cells) upon stimulation. In order to address this question, splenocytes of infected WT and LTβR^−/−^ mice (day 7 and 10 *p.i.*) were prepared and were restimulated *ex vivo* with toxoplasma lysate antigen (TLA) before flow cytometry analysis (Fig. 6).

**Fig. 6.**
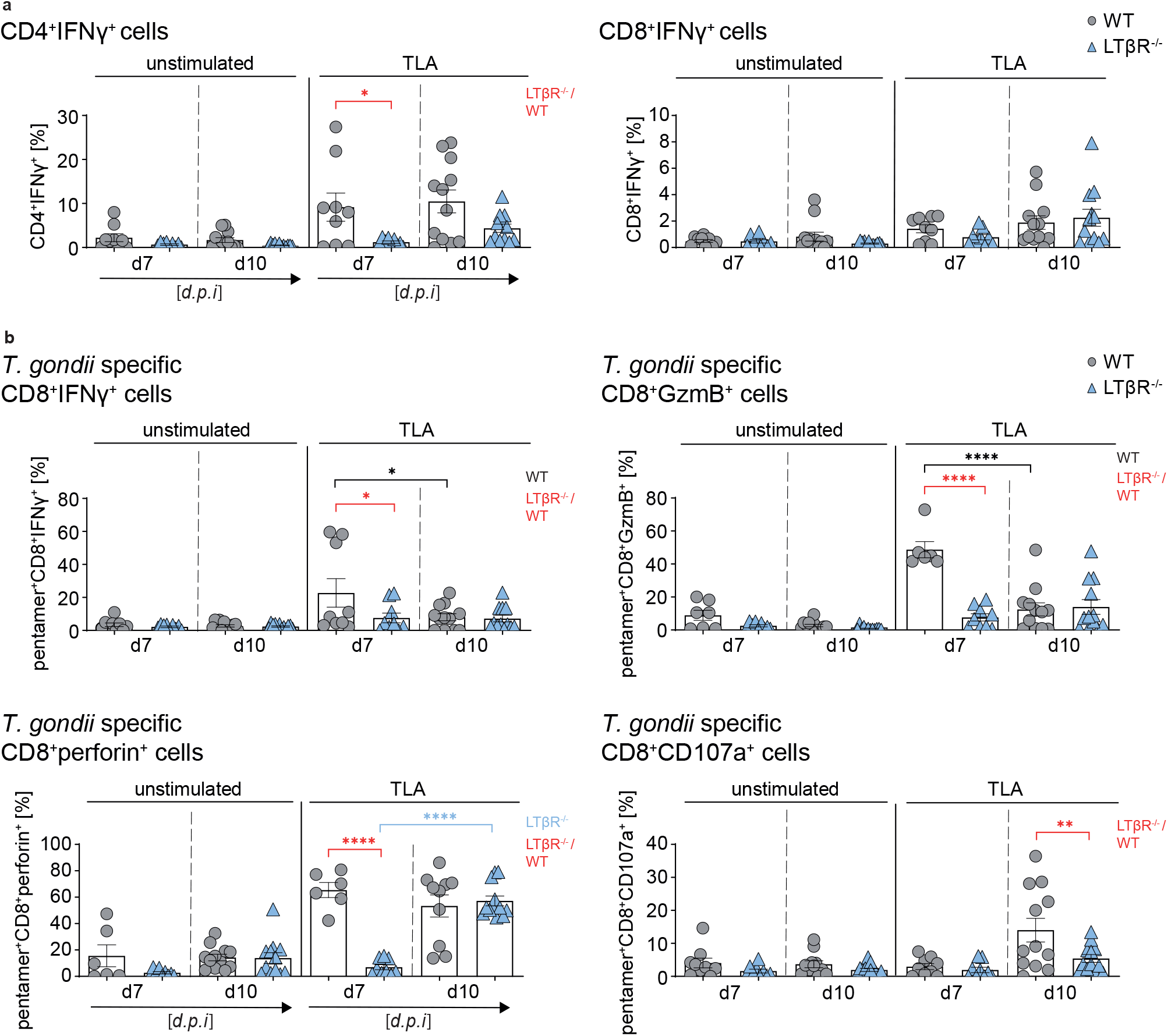
LTβR deficiency impairs T cell effector function in the spleen. Intracellular staining of **a**, CD4^^+^^IFNγ^+^ and CD8^+^IFNγ^+^ T cells [%] and **b**, cytotoxic granule (GzmB^+^ or perforin^+^) containing and degranulating (CD107a^+^) pentamer^+^ CD8^+^ T cells of unstimulated and toxoplasma lysate antigen (TLA) *ex vivo* restimulated splenocytes from *T. gondii* infected (d7 and 10 *p.i.*) WT and LTβR^−/−^ mice (d7: n≥6, d10: n≥10). Representative data of at least two independent experiments; symbols represent individual animals, columns represent mean values and error bars represent ± SEM. 2way ANOVA corrected for multiple comparison by the Tukey‘s post hoc test was used for statistical analysis. *P<0.0332, **P<0.0021, ****P<0.0001.

After *ex vivo* TLA restimulation LTβR^−/−^ T cells compared to WT T cells showed a significantly reduced frequency of CD4^+^ IFNγ producing T cells in splenocytes from day 7 *p.i.* and a reduced percentage in splenocytes at day 10 *p.i.* Similar frequencies for CD8^+^ IFNγ producing T cells could be detected in restimulated splenocytes for both genotypes on both days (Fig. 6a). There were no significant differences between the two genotypes for granzyme B containing CD8^+^ cells in restimulated cells from either day 7 or day 10 *p.i.* (Suppl. Fig. 6a). For CD8^+^perforin^+^ cells, WT mice showed higher frequencies in day 10 restimulated cells, but LTβR^−/−^ mice showed a delayed but significant increase from day 7 to day 10 resulting in frequencies similar to those of WT mice for day 10 *p.i.* (Suppl. Fig. 6a). In contrast, CD8^+^107a^+^ T cell frequencies in WT spleens increased significantly in restimulated cells from day 10 *p.i.* and were significantly higher than that of LTβR^−/−^ spleens (Suppl. Fig. 6a). When we directly analyzed IFNγ^+^GzmB^+^ and IFNγ^+^perforin^+^ cells, we found a significantly higher frequency in restimulated splenocytes of WT mice at day 7 *p.i.* compared to LTβR^−/−^ mice (Suppl. Fig. 6b).

For *T. gondii* specific (pentamer^+^) CD8^+^ T cells (Fig. 6b) we found a significantly higher frequency of pentamer^+^CD8^+^ IFNγ producing T cells in restimulated splenocytes from WT mice on day 7 *p.i.* compared to LTβR^−/−^ mice but not in restimulated splenocytes from day 10 *p.i.* Interestingly, *T. gondii* specific CD8^+^GzmB^+^ T cells showed a similar picture: A significantly increased frequency in restimulated splenocytes from WT mice on day 7 *p.i.* as compared to LTβR^−/−^ mice, and no difference of these cells in splenocytes from day 10 *p.i.* In WT compared to LTβR^−/−^ spleens, *T. gondii* specific CD8^+^perforin^+^ T cells were also significantly higher in restimulated WT splenocytes at day 7 *p.i.* However, here, LTβR mice showed a significantly increased frequency of CD8^+^perforin^+^ T cells in restimulated splenocytes from day 10 compared to day 7 *p.i.* resulting in similar frequencies for WT and LTβR^−/−^ CD8^+^perforin^+^ T cells at day 10 *p.i.* Finally, the percentage of *T. gondii* specific CD8^+^CD107a^+^ T cells was similar for both genotypes in restimulated splenocytes at day 7 *p.i.*, but significantly increased for WT mice in restimulated splenocytes from day 10 *p.i.* whereas only few CD8^+^ LTβR^−/−^ T cells degranulated. To summarize, importantly, parasite specific granzyme B granule containing (pentamer^+^CD8^+^GzmB^+^) as well as degranulating (pentamer^+^CD8^+^CD107a^+^) T cells do not appear to be detectable in LTβR^−/−^ mice after *T. gondii* infection, whereas the increase of parasite specific perforin granule containing (pentamer^+^CD8^+^perforin^+^) T cells seems to be delayed in LTβR^−/−^ than WT mice. These results demonstrate that while the T cell compartment does not seem to be affected in regard to cell numbers LTβR^−/−^ mice show a clear functional defect in the parasite specific CD8^+^ T cell compartment as well as clearly decreased IFNγ producing CD4^+^ T cells after infection.

### LTβR deficiency abrogates *T. gondii* specific isotype class switching

RNAseq data of lung tissue from uninfected (day 0) and *T. gondii* infected (day 7 *p.i.*) WT and LTβR^−/−^ animals was further analyzed to elucidate the interaction between *T. gondii* and host immune responses. The data was filtered for differentially expressed genes, hierarchical clustering was performed and illustrated as sample dendrogram with a trait heat map (Suppl. Fig. 7a) for identification of possible outliers. All tested samples showed adequate clustering and could accordingly be grouped into uninfected and infected WT and LTβR^−/−^ mice. Next, gene expression data was condensed into ten module eigengenes (ME0 - ME9; Suppl. Fig. 7b) and used to generate a host-pathogen network prediction model (Fig. 7a) displaying the relationship between modules (ME) and experimental conditions. This model captures the influence of *T. gondii* infection (Infection), the LTβR^−/−^ genotype (Genotype), and total *T. gondii* genes (X) on host gene modules (ME0 - ME9) detected in each sample. Upon closer inspection, this model shows that LTβR expression (contained in ME6) is suppressed by the LTβR^−/−^ genotype, which fits our experimental conditions. This model predicts that in WT mice high expression of genes contained in ME6 suppresses genes contained in ME4 (Top GO term ‘B cell receptor signaling pathway’), while enhancing gene expression in ME3 (Top GO term ‘lymphocyte differentiation’). This implies that the loss of the LTβR slightly increases ME4 levels (Suppl. Fig. 7c; Top GO term B cell receptor signaling pathway, Fig. 7a) containing genes for immunoglobulin production and humoral immune response mediated by circulating immunoglobulin during *T. gondii* infection. Furthermore, the network predicts that in LTβR^−/−^ mice *T. gondii* infection reduces ME3 levels (Suppl. Fig. 7d; Top GO term lymphocyte differentiation, Fig. 7a) containing genes for B cell activation and isotype switching. In addition, GSEA generated from RNAseq data also showed significant upregulation of these pathways, indicating a disturbed B cell response (Suppl. Fig. 2).

**Fig. 7.**
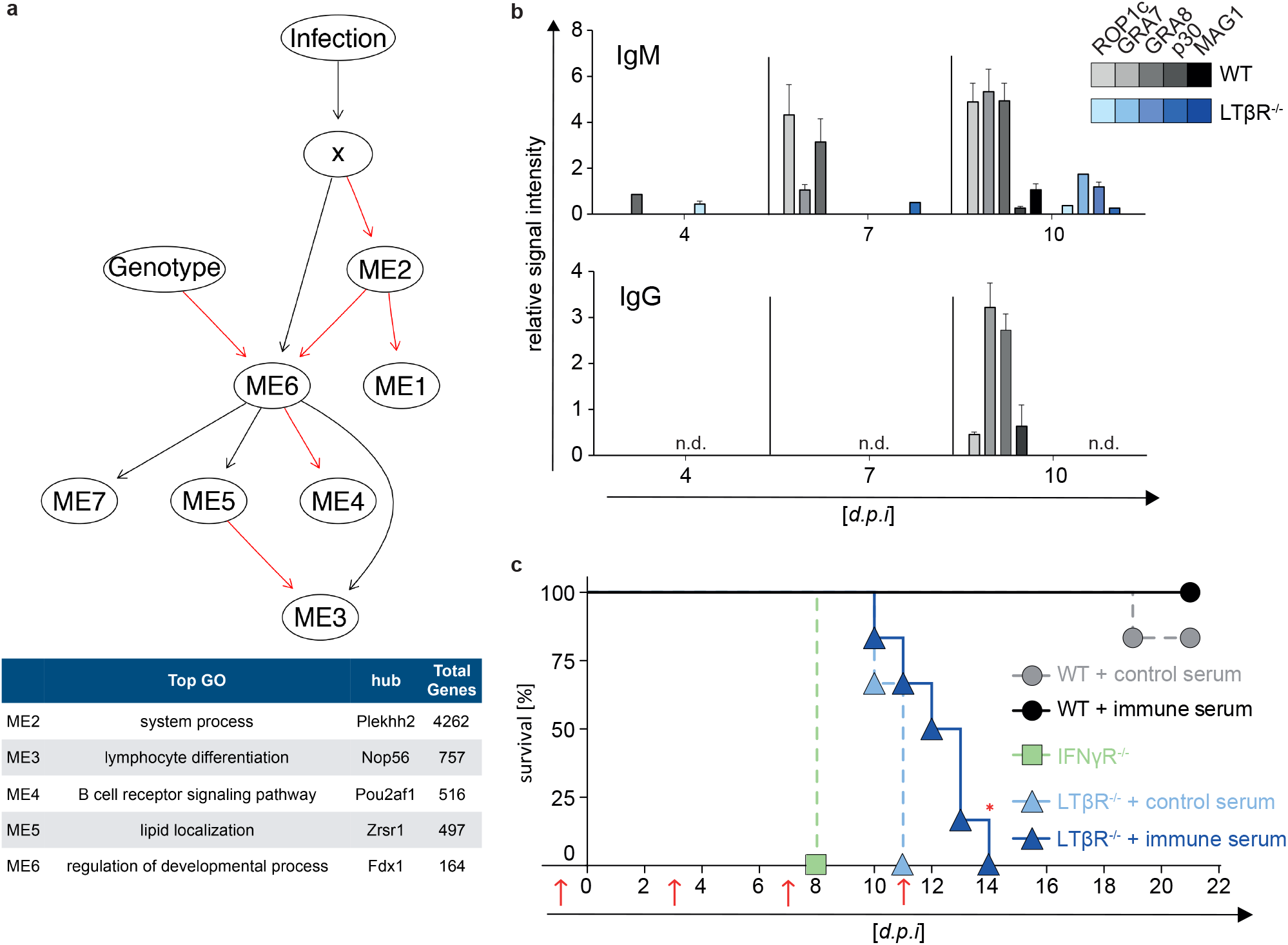
Abrogated parasite specific isotype class switching and reconstitution of mice with *T. gondii* specific immune serum. **a**, Host-pathogen network prediction model generated on RNAseq data of lung tissue of uninfected (d0) and *T. gondii* infected (ME49, 40 cysts; d7 *p.i.*) WT and LTβR^−/−^ mice (n=3/group). GmicR was used to detect relationships between module eigengenes (ME) and experimental conditions. x represents total *T. gondii* gene expression data for each sample, infection and genotype were included as variables. Red lines indicate inverse and black lines positive relationships. Representative gene ontologies and hub genes reported by GmicR for each module are shown in the summary table. **b**, *T. gondii* specific IgM and IgG antibody response in serum of uninfected (d0) and *T. gondii* infected (ME49, 40 cysts *i.p.*) WT and LTβR^−/−^ mice (d4 and d7 *p.i.*: n=15, d10 *p.i.*: n≥20). Shown is a representative result of four independent experiments, bars represent mean values ± SEM. **c**, Transfer of serum (red arrows; d-1, 3, 7 and 11 *p.i.*) from uninfected donor WT mice (control serum) or from *T. gondii* infected (ME49, 20 cysts, *i.p.*)donor WT mice (immune serum) into WT and LTβR^−/−^ acceptor mice. On day 0, acceptor mice (n=6/group) were infected with *T. gondii* (ME49, 10 cysts, *i.p.*) and survival was evaluated. IFNγR^−/−^ mice (n=3) served as infection controls. Data shown in **c** represent one experiment. A log rank (Mantel Cox) test was used for statistical analysis represented in **c.** *P<0.0332 and n.d.= not detected.

Due to this highly surprising prediction, as well as the different B cell numbers of WT and LTβR^−/−^ in the spleen on day 10 *p.i.* (Fig. 5a & c), we then asked whether an altered B cell mediated humoral immune response could be directly involved in the high mortality of LTβR^−/−^ mice after *T. gondii* infection. The presence of immunoglobulin (Ig) M and IgG antibodies specific for *T. gondii* antigens was determined during the acute phase of infection (days 4, 7, and 10 *p.i.*) using line blots coated with specific recombinant *T. gondii* tachyzoite and bradyzoite antigens (ROP1c, GRA7, GRA8, p30 and MAG1). LTβR^−/−^ mice compared to WT mice showed a delayed and reduced *T. gondii* specific IgM and, suprisingly, an abrogated *T. gondii* specific IgG antibody response in the serum during infection (day 4, 7, and 10 *p.i.;* Fig. 7b), demonstrating a lack of functional isotype switching that is in line with the bioinformatic host-pathogen prediction network.

### LTβR deficiency can be partially compensated for by transfer of *T. gondii* immune serum

Since it has been described that a *T. gondii* specific IgG response is required for a reduction of the parasite burden (25, 51), we treated LTβR^−/−^ mice with serum from *T. gondii* infected WT animals (immune serum) and uninfected mice (control serum) and monitored survival after *T. gondii* infection (Fig. 7c). Serum transfer experiments showed that LTβR^−/−^ mice treated with immune serum exhibit significantly prolonged survival (up to day 14 *p.i.*) compared to littermates that received control serum which died by day 11 *p.i*. IFNγR^−/−^ mice served as infection control and succumbed as reported around day 8 *p.i.* (53). These data demonstrate that LTβR mediated signaling is essential for the development of an efficient humoral immune response to *T. gondii* infection.

## Discussion

The results obtained in this study corroborate a profoundly deficient immune response of LTβR^−/−^ mice to *T. gondii* infection and reveal an impaired IFN response, a severe functional T cell defect as well as a humoral immune deficiency in the absence of LTβR.

One reason for the significantly increased parasite burden and significantly reduced survival rates of LTβR^−/−^ mice is the inadequate cytokine, especially the IFNγ, response. The elevated levels of LTα and significantly increased levels of LTβ in the lung of LTβR^−/−^ mice could be caused by compensatory mechanisms and/or lack of negative feedback mechanisms due to the absence of the LTβR. Since we also found elevated levels for IFNγ, IL-6, IFNβ, IL1α, IL-17A, significantly elevated expression levels for IL-1β in the serum and for IL-4 in the lung, we suggest that overall, uninfected LTβR^−/−^ mice show a dysregulated, more activated, albeit stable immune homeostasis. This is in accordance with the finding that LTβR^−/−^ animals present with splenomegaly, most probably due to microbiota-mediated inflammation (54). When *T. gondii* infection disrupts this precarious balance in LTβR^−/−^ mice the dysregulation becomes more pronounced: On the one hand, LTβR^−/−^ mice have lower levels of IFNγ in the serum early during infection, but on day 10 *p.i.* when WT mice already show decreased IFNγ levels, they remain high in LTβR^−/−^ mice, not only in serum but also in the lungs. Conversely, IL-6 expression in the serum is markedly increased in LTβR^−/−^ mice compared to WT mice throughout the infection. This suggests that by day 10 *p.i.* parasite expansion is being controlled in WT but not LTβR^−/−^ mice. The significantly increased levels of IL-10 in LTβR mice on day 10 *p.i.* could be a protective/counteractive mechanism to prevent extensive immunopathology (55). Interestingly, several cytokines in LTβR^−/−^ mice are transiently but significantly upregulated on day 4 *p.i.* This also suggests a disruption of the precarious immune homeostasis in LTβR mice. In contrast to the activated immune homeostasis in LTβR^−/−^ mice they show decreased expression levels for chemokines/chemokine receptors, genes involved in IFNγ signaling and IFNγ induced genes in the lung on day 7 *p.i.* This points towards an inability of LTβR^−/−^ mice to mount an efficient immune response to *T. gondii* infection and is supported by the finding that upregulation of IFNγ regulated effector molecules known to be important for *T. gondii* containment such as iNOS, IDO1, NOX2-gp91phox and mGBPs is deficient in LTβR animals. In the lung, in the case of NOX2-gp91phox, this could be due to the lack of TNFα expression, as it has been shown in ex vivo experiments for Bronchoalveolar Fluid Cells and Human Pulmonary Artery Endothelial Cells that TNFα upregulates NOX2-g91phox (56, 57). The mRNA expression profile of mGBPs and protein expression of mGBPs 2 and 7 also fits into this pattern: mGBPs are essential for efficient control of *T. gondii* expansion (31, 33, 58) and RNAseq analysis shows that uninfected LTβR mice have overall increased expression of mGBPs while infected animals show overall less upregulation. And while LTβR^−/−^ animals do upregulate mGBP expression during the course of infection, they show significantly lower expression on day 10 *p.i.* compared to WT mice in all cases except for mGBP6/10.

LTβR^−/−^ animals also show increased baseline expression of IFNγ mRNA in the lung, which would explain the elevated baseline JAK1 protein expression. Increased JAK expression should lead to increased JAK phosphorylation and consequently increased STAT1 recruitment and STAT1 phosphorylation (59, 60). However, we observed delayed upregulation of STAT1 and less pSTAT1 protein in infected LTβR^−/−^ animals and therefore hypothesize that the lack of LTβR signaling somehow affects STAT1 expression or recruitment via a so far unknown mechanism. Notably, Kutsch et al. also showed reduced STAT1 expression in LTβR^−/−^ mice (61).

We conclude that, due to the underlying dysregulation of the immune homeostasis, LTβR^−/−^ mice are unable to initiate a coordinated immune response leading to either delayed upregulation of essential cytokines (e.g. IFNγ) or over-expression of others (e.g. IL-6, TNF). This is also supported by our findings, that LTβR^−/−^ mice do not show the typical splenomegaly associated with (*T. gondii*) infection (39).

In line with published data, we found a virtual absence of NK1.1^+^ cells in LTβR^−/−^ mice, and NK1.1^+^ cell numbers dropped in WT mice after infection, most probably due to conversion into ILCs (51). Also, a lack of NKT cells has been shown for LTβR^−/−^ mice (62). Interestingly, a dual role for NKT cells in *T. gondii* infection has been described: On the one hand, they are able to release large amounts of IL-4 and IFNγ upon activation and to shift the T cell response towards a Th1 pattern, and on the other hand, the uncontrolled Th1 response can lead to severe immunopathology (63). Since they have also been indicated in the suppression of a protective immunity against *T. gondii* infection (64) it is maybe not surprising that their numbers are downregulated after infection in WT animals.

Overall, we did not find T cell numbers to be significantly different in either uninfected or *T. gondii* infected LTβR^−/−^ mice compared to WT mice. However, we found profound defects in T cell effector functions: The reduced number of IFNγ producing CD4^+^ T cells and functional *T. gondii* specific CD8^+^ cytotoxic lymphocytes (GzmB^+^, perforin+, CD107a^+^) strongly implies that cytotoxic T cell mediated killing is severely impaired in LTβR^−/−^ animals. Since these responses are known to be essential for efficient *T. gondii* containment, this marked functional deficiency is probably one reason for the susceptibility of LTβR^−/−^ mice to the parasite.

In contrast to T cell numbers, B cell numbers differed significantly in LTβR^−/−^ mice compared to WT mice. On day 10 post infection, numbers of CD19^+^ B cells in WT spleens were significantly lower compared to those of LTβR^−/−^ animals. This is most probably due to maturation of B cells to IgG producing plasma cells in WT mice, which emigrate to the bone marrow and lose surface CD19 in the process. In LTβR^−/−^ mice the lack of class switching would inhibit maturation and migration of B cells to the bone marrow.

Since the host-pathogen network prediction model we generated from *T. gondii* infected mice indicated that the loss of the LTβR inhibits B cell responses including isotype switching in *T. gondii* infection we further analyzed the humoral immune response. We were able demonstrate that *T. gondii* infected LTβR^−/−^ mice produced less *T. gondii* specific IgM compared to WT mice, and no detectable *T. gondii* specific IgG. In as much this failure is due to impaired IFNγ production which is an important cytokine for isotype class switching (65) will be determined in the future.

While Glatman Zaretzky et al. (39) argue that the disrupted lymphoid structure, which includes the lack of defined germinal centers in LTβR^−/−^ mice is the main cause of the reduced antibody response, Ehlers et al. (5) show via BM chimeras that the effects of LTβR deficiency in *M. tuberculosis* infection cannot be attributed solely to the architectural differences, but are also directly caused by the lack of LTβR mediated signaling. LTα, another member of the TNF/TNFR superfamily, has similar but not identical functions to LTβ in the development of secondary lymphoid organs and immune modulation (2). LTα^−/−^ animals also present with a disturbed architecture of the lymphoid system (no LNs, no PPs, no GCs and a disorganized white pulp) (2, 66). *T. gondii* infected LTα^−/−^ mice are shown to have reduced numbers of *T. gondii* specific IFNγ producing T cells and lower *T. gondii* specific antibody titers but BM chimera experiments demonstrated that an intact secondary lymphoid system is not sufficient to generate an effective immune response (25).

Although protective B cell responses have been described to play a more significant role in chronic rather than acute *T. gondii* infection in some *T. gondii* infection models (49–51, 67) our data indicate that a robust humoral immune response is dependent on LTβR signaling and also is a prerequisite for survival during acute *T. gondii* infection. This conclusion is validated by our data showing that the survival of LTβR^−/−^ animals can be significantly prolonged by transfer of immune serum containing *T. gondii* specific antibodies.

Finally, the host-pathogen prediction network generated in this study indicates that *T. gondii* infection suppresses B cell responses in WT animals. This could point towards an unknown *T. gondii* strategy to evade the host immune system. Early *T. gondii* mediated suppression of B cell responses could support dissemination and cyst formation in the brain, facilitating the establishment of chronic infection (25, 68). Since *T. gondii* is known to have developed different mechanisms to evade host immune responses (69), it is worth exploring this approach in the future.

Taken together, we demonstrate that the loss of LTβR signaling results in a combined and profoundly depressed IFNγ response, impaired T cell functionality and the failure to induce parasite specific IgG antibodies leading to an increase in parasite burden and fatal outcome of *T. gondii* infection. Therefore, for the first time, we suggest an LTβR mediated modulation of the IFNγ signaling pathway *in vivo*. Further understanding of this complex interplay between LTβR and IFNγ signaling pathways will provide new insights into the pathogenesis of *T. gondii* and may provide novel therapeutic strategies.

## Acknowledgments

We thank Nicole Küpper, Julia Mock and Karin Buchholz for technical assistance. This work was supported by the Jürgen Manchot Foundation (Molecules of Infection III – MOI III). Computational support of the Zentrum für Informations- und Medientechnologie, especially the HPC team (High Performance Computing) at the Heinrich Heine University is acknowledged.

## Author contributions

A.T. performed and analyzed all experiments, except for Fig. 2a, Fig. 3a, Fig. 4, Fig. 7a, supplementary Fig. 2 and Fig. 7; RVS developed the immune network models for analysis of RNAseq datasets. M.H. performed experiments illustrated in Fig. 4. P.P. and K.K. performed RNA sequencing. A.T., U.R.S. and K.P. wrote the manuscript with input from D.D., I.R.D. and C.F.W; K.P., U.R.S. and A.T. designed the study.

## Additional information

Supplementary information is available for this paper.

## Methods

### Mice

LTβR^−/−^ mice were previously described (7) and are back crossed for at least 10 generations onto a C57BL/6N background. Wild-type (WT) littermates were used as controls. Mice were kept under specific pathogen-free conditions (SPF) in the animal facility at the Heinrich Heine University Düsseldorf and were 8-16 weeks old for experiments. Cysts of the ME49 strain (substrain 2017) of *T. gondii* were collected from the brain tissue of chronically infected CD1 mice. All animal experiments were conducted in strict accordance with the German Animal Welfare Act. The protocols were approved by the local authorities (Permit# 84-02.04.2013.A495, 81-02.04.2018.A406 and 81-02.05.40.18.082). All applicable international, national, and institutional guidelines for the care and use of animals were followed.

### *Toxoplasma gondii* infection experiments

Mice were intraperitoneally infected with 40 cysts (ME49 strain) and weighed and scored daily for the duration of the experiments. Mice were euthanized on days 4, 7 and 10 post infection (*d.p.i*), uninfected mice (d0) served as controls. After euthanasia (100 mg/kg Ketamin, 10 mg/kg Xylazin, Vétoquinol GmbH) blood was taken from the Vena cava inferior and spleen, lung and muscle tissue was harvested for analysis.

### Detection of parasite load

Total DNA was isolated from tissues using a DNA isolation kit (Genekam) according to the manufacturer’s protocol. qRT-PCR was performed on a Bio-Rad CFX-96 Touch-Real-Time Detection System. TgB1 primers and probe (Metabion) were used to amplify a defined section of the 35-fold repetitive B1 gene from *T. gondii* and are listed in Supplementary Table 1. The *T. gondii* standard curve was used to determine B1 amplification for calculation of parasite load.

### Cytokine measurement

Cytokines CCL2, IFNγ, IFNβ, IL-1α, IL-1β, IL-6, IL-10, IL12p70, IL-17A, IL-23, IL-27, and TNFα were measured using the LEGENDplex™ Mouse Inflammation Panel (BioLegend^®^) according to the manufacturer’s protocol. Samples were measured using a BD FACSCanto™ II.

### Real-time qRT-PCR

Total RNA was isolated from tissues using the TRIzol reagent (Invitrogen) according to the manufacturer’s protocol. cDNA was reversely transcribed using SuperScript III reverse transcriptase (200 U/µl; Invitrogen). qRT-PCR was performed on the Bio-Rad CFX-96 Touch-Real-Time Detection System. Primer sequences and corresponding probes (Metabion, Roche & TipMolBIOL) are listed in Supplementary Table 1. Results are expressed relative to expression in untreated WT mice normalized to β*-actin* (2^−ΔΔCT^).

### RNAseq analysis

Lung tissue of uninfected (d0) und *T. gondii* infected (ME49 strain, 40 cysts*, i.p.*) WT and LTβR^−/−^ mice was obtained and RNA sequencing was performed on a HiSeq3000 device. Mouse and *T. gondii* transcripts were quantified from fastq files using Salmon with default settings and GCbias compensation. For transcriptome models, Mus musculus GRCm38 cDNA (ensembl.org, release-97) and *Tgondii*ME49 Annotated Transcripts (toxodb.org, ToxoDB-45) were used. Mouse transcripts from pseudogenes or with retained introns were excluded prior to conversion to gene counts by the DESeq2 package. Non-protein encoding *T. gondii* transcripts were excluded prior to conversion to gene counts. DEseq2 was used to test for Genotype-specific responsiveness to infection with the following model: ~ Genotype * Infection. To calculate WT-specific responsiveness, we used the following model: ~ Genotype + Genotype: Infection. For significance the Wald test with an adjusted p-value of 0.1 was used.

### Host-pathogen network generation

Previously developed analytic tools for ‘omics datasets were used to generate the host-pathogen network as described (70). Prior to network generation, the VST-normalized data were filtered for genes that showed significant differential expression for at least one contrast. This produced an expression matrix for 10,748 genes. The GmicR package was then used for module detection, using a minimum module size of 30, mergeCutHeight of 0.3, and Rsquared cut of 0.80. To detect relationships between modules and infection, VST-normalized data *T. gondii* expression levels for each sample were aggregated by sum and then this numeric data was merged to module eigengenes using the Data_Prep function of GmicR [Supplementary Figure 6]. Genotype and infection conditions were merged with the discretized data. A white list indicating the parent to child relationship from “Genotype” to “ME6” corresponding to the module containing LTβR was included in the Bayesian network learning process. A final network was generated using the bn_tabu_gen function with 500 bootstrap replicates, “bds” score, and iss set to 1. Inverse relationships between nodes were detected using the InverseARCs function from GmicR with default settings.

### Immunoblot analysis and antibodies

Tissues were homogenized in PBS containing cOmplete™ Protease Inhibitor Cocktail (Roche) using the Precellys^^®^^ homogenizer (Bertin). Protein concentration was measured using the Pierce BCA Protein Assay Kit (Thermo Scientific™) according to the manufacturer’s protocol. Samples [10 µg/lane] were separated by 4-12% SDS-PAGE, followed by electrophoretic transfer to nitrocellulose membranes before blocking and incubation with primary antibodies listed in Supplementary Table 2. HRP-labeled anti-rabbit or anti-mouse antibodies (Cell Signaling Technologies) were used as secondary antibodies. Relative signal intensity of protein bands was quantified using ImageJ (NIH).

### tSNE

The cloud-based platform Cytobank^^®^^ (71) (Mountain View) was used for visualization of flow cytometry data. 60,000 events per sample were analyzed (parameters: iterations 2,400, perplexity 80, Theta 0.5) before overlaid dot plots were generated.

### Flow cytometry

Spleens were harvested and digested 30 min. at 37 °C using Collagenase D (100 mg/ml) and DNase I (20,000 U/ml). Tissue digest was stopped using 1x PBS containing 10 mM EDTA before cell solution was filtered using a 70 μm cell strainer. A RBC lysis (Merck) was performed before cell numbers were calculated. Single-cell suspended splenocytes (1×10^6^ cells) were stained with the Fixable Viability Dye eFluor^®^ 780 (eBioscience™). Surface staining with antibodies specific for CD3e (145-2c11), CD4 (RM4-5), CD8a (53-6.7), CD19 (6D5), CD25 (3C7), and NK1.1 (PK136) all purchased from BioLegend (expect for CD4 purchased from BD Bioscience), was performed. For intracellular staining splenocytes were incubated for 20 h with toxoplasma lysate antigen (TLA, 15 μg/ml) before adding brefeldin A (eBioscience™) for an additional 4 hours. After surface staining with anti-CD4 (RM4-5), anti-CD8a (53-6.7), anti-CD107a (1D4B), and anti-TCRb (H57-597) cells were fixed, permeabilized and stained with anti-IFNγ (XMG1.2), anti-granzyme B (QA16A02), and anti-perforin (S16009A) (all purchased from BioLegend) using Fix & Perm^®^ Cell Permeabilization Kit (Life Technologies) according to the manufacturer’s protocol. Major histocompatibility complex class I - SVLAFRRL pentamer was purchased from ProImmune and used in experiments as indicated. BD Calibrate beads (BD Bioscience) were added to the samples before acquisition with a BD LSRFortessa.

### Detection of *T. gondii* specific antibodies

*Recom*Line *Toxoplasma* IgG/IgM kit (Mikrogen Diagnostik) was used to detect IgM and IgG antibodies against *T. gondii* in serum. Anti-human IgM and IgG conjugates provided within the kit were replaced with anti-mouse IgM-HRP-labeled (Invitrogen) and anti-mouse IgG-HRP-labeled (Invitrogen) conjugates. Otherwise, the assay was performed according to the manufacturer’s protocol.

### Serum transfer

Blood from naïve donor mice (control serum) or WT mice infected with 20 cysts *i.p.* of the ME49 strain of *T. gondii* (immune serum) was collected from the vena cava inferior. After 2h incubation at RT serum was collected by centrifugation of the blood. Acceptor WT and LTβR^−/−^ mice were reconstituted intraperitoneally with 0.2 ml serum one day prior to infection (d-1) as well as on days 3, 7 and 11 *p.i.* Acceptor (WT and LTβR^−/−^ mice) as well as IFNγ^−/−^ control mice were intraperitoneally infected with 10 cysts (ME49 strain) and weighed and scored daily for the duration of the experiment. *T. gondii* specific antibodies were detected via Line Blots to confirm the presence and assess the amount of *T. gondii* specific antibodies in control and immune serum.

### Statistical analysis

Data were analyzed with Prism (Version8, GraphPad) using log rank (Mantel Cox) test or 2way ANOVA corrected for multiple comparison by the Tukey’s or Sidak’s post hoc test as indicated in the figure legends. Symbols represent individual animals, columns represent mean values and error bars represent ± SEM. P values of ≤0.0332 were considered statistically significant and marked with asterisks. P values of ≥0.0332 were considered statistically not significant and were not specifically marked.

## Data availability

The data that support the findings of this study are available from the corresponding author.

